# Reduced Medial Frontal Positivity During the Stimulus-Response Interval Precedes Action Errors and Explains Task Deficits in Attention-Deficit Hyperactivity Disorder

**DOI:** 10.1101/412494

**Authors:** Scott J. Burwell, Scott Makeig, William G. Iacono, Stephen M. Malone

## Abstract

Brain mechanisms responsible for errors during cognitive tasks are poorly understood, particularly in adolescents with attention-deficit hyperactivity disorder (ADHD). Using subject-specific multimodal imaging (EEG, MRI, behavior) during flanker task performance by a sample of 94 human adolescents (mean age = 15.5 years, 50% female) with varying degrees of ADHD symptomatology, we examined the degree to which amplitudes of source-resolved event-related potentials (ERPs) from brain independent components within a critical (but often ignored) period in the action selection process, the stimulus-response interval, predicted motor response errors (across trials) and error rates (across individuals). Reduced amplitudes of Frontocentral P3 (peaking at approximately 390 milliseconds in stimulus-locked ERPs) and Pre-Movement Positivity (PMP, peaking at approximately 110 milliseconds pre-response in response-locked ERPs) in projections from posterior medial frontal cortex (pMFC) predicted erroneous responses, and reduced amplitude of PMP predicted a larger participant error rate. After regressing stimulus-from response-locked ERPs, we concluded that errors primarily depended upon response selection processes reflected in PMP amplitude. Finally, mediation analyses showed that smaller PMPs on correct response trials was associated with the higher frequency of errors committed by adolescents with more ADHD symptoms. These results bolster the importance of pMFC in action selection and support the possible value of using PMP as an intervention target to remediate performance deficits in ADHD.

## Introduction

Brain mechanisms responsible for errors during cognitive tasks remain elusive, such that action monitoring research has focused on brain physiology that occurs too early (preceding the error trial by one or more trials) or too late (following the error) to enable insight into perceptual and motor processes occurring during the error trial itself. Yet, such insights may be valuable for understanding why individuals with attention-deficit hyperactivity disorder (ADHD) make many mistakes during cognitive tasks (Lijffijt et al., 2005; Mullane et al., 2009) and are also more accident prone in myriad life situations (Swensen et al., 2004; Brook and Boaz, 2006; Jerome et al., 2006; Vaa, 2014).

Identifying functional brain antecedents to errors is important for understanding possible causes of error proneness in ADHD. Studies of healthy controls have revealed that errors may be foreshadowed by hemodynamic deactivations (Weissman et al., 2006; Eichele et al., 2008) and event-related potential (ERP) amplitude reductions (Ridderinkhof et al., 2003; Allain et al., 2004; Hajcak et al., 2005; O’Connell et al., 2009; Masaki et al., 2012) in frontal cortex one to several trials before the error trial. Similar processes have been posited to explain attentional lapses in ADHD that may endure 10 or more seconds (Sonuga-Barke and Castellanos, 2007; Yordanova et al., 2011), but this line of research gives little insight into brain physiology on individual error trials, specifically within the brief action selection period (≈ 0.5 second) between stimulus presentation and the motor response.

Aberrant neurophysiology during the *stimulus-response interval (SRI)* on speeded motor response tasks may enable insight into erroneous action selections. For example, larger positive peaks occurring approximately 300 milliseconds in stimulus-locked ERPs over frontal cortex have correlated with faster motor responses (Makeig et al., 1999; Delorme et al., 2007) and better accuracy (Perri et al., 2014; Perri et al., 2015). Additionally, response-locked ERP amplitudes preceding manual responses have been linked to performance errors (Meckler et al., 2013; Bode and Stahl, 2014; Roger et al., 2014). Yet, error associations during the SRI remain difficult to interpret because stimulus- and response-locked ERPs are often studied separately, and the rapid succession among stimulus and response waveforms suggests “temporal confounding” (Smith and Kutas, 2015a, b), meaning it is unclear whether error-related effects during the SRI should be attributed to stimulus-evoked (perceptual) or response-preceding (motor) processing.

Such methodological limitations contribute to the gap in knowledge regarding SRI error-related brain dynamics in ADHD. While studies using functional magnetic resonance imaging (fMRI) have found frontal brain hemodynamic differences in ADHD during action monitoring (e.g., Suskauer et al., 2008; Chao et al., 2009), the timings of such effects are unclear due to fMRI’s poor temporal resolution (Cohen, 2011). Also, ERP studies on ADHD subjects have largely ignored error and correct trial differences during the SRI, either focusing on stimulus-locked ERPs solely in correct trials, or on response-locked ERPs solely following errors (e.g., Albrecht et al., 2008; McLoughlin et al., 2009; Karch et al., 2010; Yoon et al., 2013; Burwell et al., 2014).

Here, we sought to identify stimulus- and response-locked brain potentials occurring within the SRI (between visual stimulus presentation and the button response) that are associated with errors (within individuals) and error rates (across individuals) during a speeded response “flanker” task (Eriksen and Eriksen, 1974) in a sample of adolescents with varying degrees of ADHD symptomatology. We limited our investigation to effective brain sources using electroencephalogram (EEG) independent component analysis decomposition and equivalent current dipole localization within models constructed from subjects’ MRI head images. Additionally, to disambiguate temporal confounding among overlapping stimulus-versus response-related processes, we introduced a novel regression-ERP approach akin to methods used to disentangle event-related fMRI responses. We predicted that brain potentials found to be related to errors would themselves predict the increased error rate of individuals having more ADHD symptoms.

## Materials and Methods

### Participants

Forty-eight pairs of monozygotic twins (*M* [*SD*] age = 15.5 years [.9], 50% female) were enrolled in the AdBrain study (for more information on this study, see: Malone et al., 2014; Silverman et al., 2014; Wilson et al., 2015; Burwell et al., 2016) conducted under the auspices of the Minnesota Center for Twin and Family Research (MCTFR) at the University of Minnesota (UMN). Diagnostic interviews and EEG recording took place at the MCTFR; neuroimaging took place on a separate day at UMN’s Center for Magnetic Resonance Research (CMRR). On average, assessments at the MCTFR and CMRR were separated by 10.6 days, with 83.3% of subjects completing both within two weeks.

Lifetime symptoms of ADHD were assessed using the Diagnostic Interview for Children and Adolescents (DICA, parent and child versions; Reich, 2000), modified to include *Diagnostic and Statistical Manual of Mental Disorders* (4^th^ ed., text revision, *DSM*; American Psychiatric Association, 2000). Symptoms were assigned if endorsed by either parent or adolescent (Leckman et al., 1982; Kosten and Rounsaville, 1992). The presence of diagnosable ADHD in this community sample was substantial (12.5%) and the number of ADHD symptoms per subject (*M* [*SD*] = 2.6 [3.9]) was elevated.

### Electrophysiological assessment

#### Flanker task

Subjects were seated in a comfortable chair in a sound attenuated, dimly lit room. An index finger response button was positioned on each armrest. Subjects performed a modified version of the flanker task (Eriksen and Eriksen, 1974). Task stimuli consisted of five-character arrays of the letters *S* and *H*. Four such target arrays appeared on screen in pseudo-random order with the following relative frequencies: *SSSSS* (33.3%); *HHHHH* (33.3%); *SSHSS* (16.7%); and *HHSHH* (16.7%). In each array, the target stimulus was the central character, which determined the desired subject response (button press); the other four characters (*S* or *H* “flankers”) had no relevance to the intended response. On each trial, the target and flanker characters were presented simultaneously at the center of a computer screen facing the subject. Before the task training block (*N* trials = 20), subjects were instructed to indicate by pressing the left-button that the target stimulus was an *S,* and to indicate with a right-button press that the target stimulus was an *H*, and that these hand-to-letter assignments would vary across subsequent task blocks. Each array was presented for 100 milliseconds. Stimulus presentations were separated by a mean of 2190 milliseconds (*SD* = 60), varying pseudo-randomly across trials in the range 2000 to 2300 milliseconds. Valid responses were only recorded when the subject responded within 1150 milliseconds following stimulus onset. Subjects were instructed to respond as quickly and accurately as possible; we checked their performance in the training block prior to beginning the EEG recording to ensure that they understood the task.

EEG was recorded during three blocks of 150 trials. Target-stimulus hand mapping was alternated in successive blocks: Block 1 (*S* = right, *H* = left), Block 2 (*S* = left, *H* = right), and Block 3 (*S* = right, *H* = left). Trials in which central and flanker stimuli were the same (e.g., central = *S*, flanker = *S*) were termed “congruent” stimulus trials; trials in which central and flanker stimuli were not the same (e.g., central = *S*, flanker = *H*) were termed “incongruent” stimulus trials.

Data for 10 task blocks in which response accuracy fell below chance (e.g., because of incorrect hand mapping during the block) were discarded. Following this process, one subject was excluded because his overall error rate (percentage of incorrectly performed responses) exceeded 40%. Two additional subjects were excluded because EEG data were contaminated with non-brain artifact. The remaining 93 subjects possessed a mean error rate of 4.9% (*SD* = 4.5) and 8.0% (*SD* = 7.4) for congruent and incongruent trials, respectively.

#### EEG recording and processing

EEG data were recorded continuously during performance of the flanker task (61 scalp electrodes; a 10/10 System electrode cap; 1024 Hz sample rate; pass-band, DC to 205 Hz) with a BioSemi ActiveTwo system (BioSemi, Amsterdam, Netherlands). Eye-movement related contributions to the EEG signals were monitored using a pair of electrodes placed above and below the right eye and another pair of electrodes placed on left and right temples. A common recording reference was used for EEG and eye-movement channels. Custom MATLAB (The MathWorks Inc., Natick, MA) scripts using functions from the EEGLAB software environment (Delorme and Makeig, 2004) were used for removing artifact-contaminated time-segments and channels by a method previously described (Burwell et al., 2016). Artifact-cleaned data were down sampled to 256 Hz, high-pass filtered above 0.1 Hz, and re-referenced at each time point to the average potential of all the channels.

#### Subject EEG source separation, dipole localization, and inter-subject co-registration

Scalp-recorded EEG data channels contain a spatiotemporal mixture of multiple brain and non-brain source activities. To “un-mix” putative brain sources from other brain and non-brain sources whose volume-conducted potentials were summed in the scalp electrode signals, we decomposed each subject’s continuous EEG using Adaptive Mixture Independent Component Analysis (AMICA; Palmer et al., 2006; Palmer et al., 2007; Delorme et al., 2012) to obtain effective EEG source activities that exhibit maximal temporal independence (see **Figure 1A** and **B**). Low-frequency (< 1.0 Hz) drifts in EEG, in some cases reflecting sweat or electrode artifacts, often account for much variance in the data and may introduce spatiotemporal non-stationarity, which tends to adversely impact ICA decompositions (Debener et al., 2010). Therefore, AMICA models were trained on data that was high-pass filtered above 1.0 Hz. For ERP analyses the resultant ICA decomposition was applied to the 0.1-Hz high-pass filtered data (Debener et al., 2010; Winkler et al., 2015). Channel weights (*W*^−1^ in **Figure 1B**) for each effective source (independent component) were then used for subsequent dipole localization.

**Figure 1.**
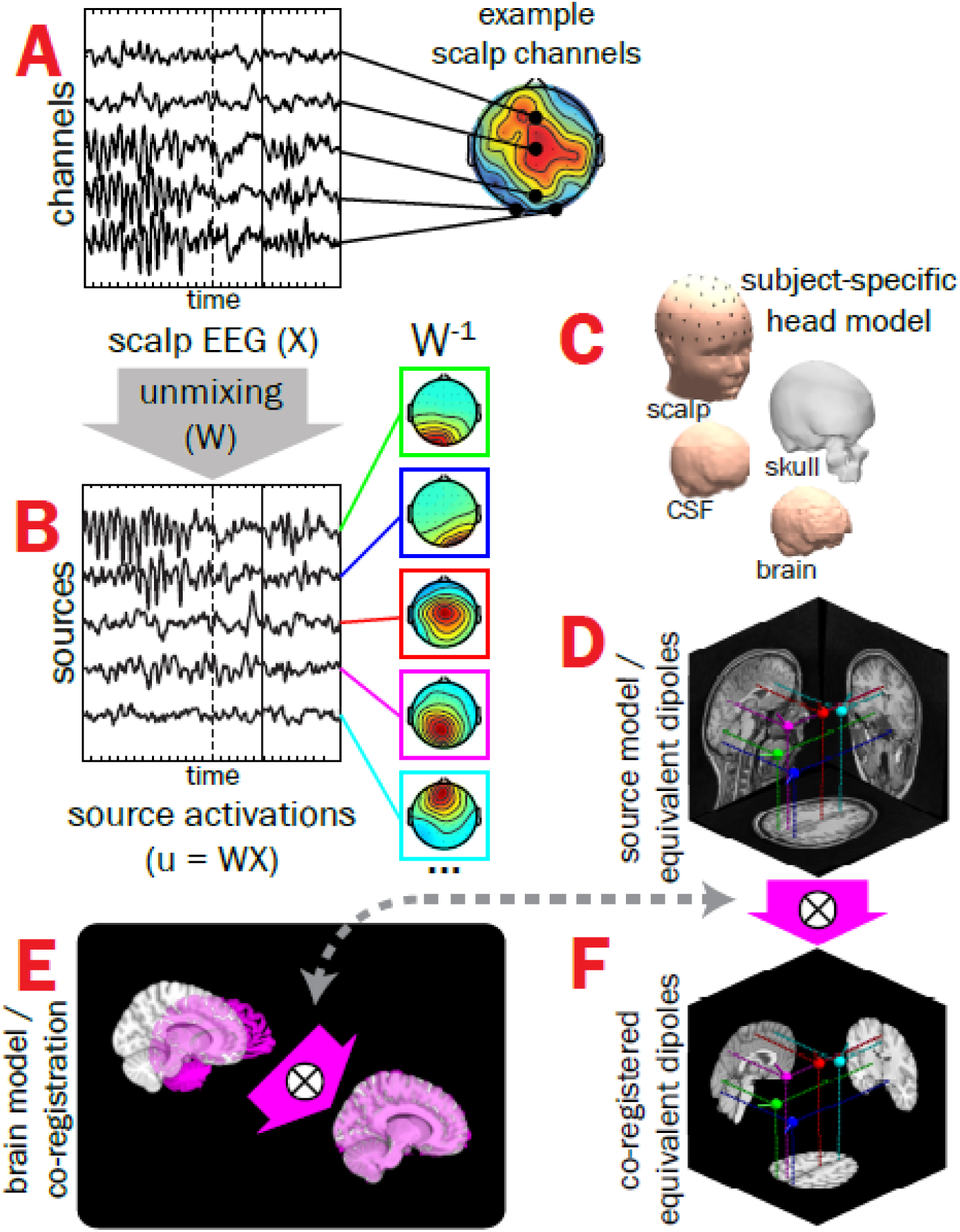
Subject EEG effective source separation, dipole localization, and inter-subject co-registration. A) Time-varying EEG recorded from a subset of scalp-positioned channels are plotted for a single subject during one correctly-performed trial of the flanker task (stimulus-onset = dashed vertical line, button-press = solid vertical line); note that scalp EEG for a given channel is a spatiotemporal admixture of activities from multiple brain and non-brain sources, leading to multiple “hot spots” in scalp projections. B) Adaptive Mixture Independent Component Analysis (AMICA) was used to “unmix” scalp-recorded channel data (by finding the un-mixing matrix, W) into putative EEG effective “source” signals with maximal temporal independence (only a subset shown here on the left) and associated scalp topographies (middle, W^-1^). C) Subject-specific head models created from structural MRIs were then used with scalp topographies to estimate the coordinates of each source’s equivalent current dipole (D). E) Finally, warping parameters mapping a given subject’s brain volume (magenta) onto the MNI template brain (gray) were obtained and applied to dipole coordinates, resulting in all subjects’ source model dipoles having a common neuroanatomical reference (F). X = scalp-recorded EEG (channels-by-time); W = unmixing matrix derived from ICA; u = EEG effective source activations (sources-by-time); W^-1^ = mixing matrix (component scalp topographies).

For subject-specific dipole localization, MRI T1-weighted anatomical images were acquired from each subject using a 3-T Tim Trio scanner (Siemens Medical Systems, Erlangen, Germany) and a magnetization prepared rapid gradient echo (MPRAGE) sequence (TR = 2530 milliseconds, TE = 3.65 ms, flip angle 7°, matrix size = 256 x 256 with a FOV of 256, 240 sagittal slices with 1 mm isomorphic voxels). Using the Neuroelectromagnetic Forward Head Modeling Toolbox (NFT; Acar and Makeig, 2010), a realistic boundary-element method (BEM) head model consisting of roughly 7,000 nodes for each layer of scalp, skull, cerebrospinal fluid (CSF), and brain mesh was generated from the anatomical image; electrode locations were spatially registered with scalp surface by aligning fiducials with nasion and two pre-auricular points (see **Figure 1C**). The forward model estimation (which maps the amplitude of each dipole in source-space to the potential at each scalp-positioned channel) used recommended conductivity values for scalp (.33 Siemens [S]/meter [m]; Geddes and Baker, 1967), skull (25:1 brain-to-skull ratio, or .0132 S/m; Lai et al., 2005; Akalin Acar and Makeig, 2013), CSF (1.79 S/m; Baumann et al., 1997), and brain (.33 S/m; Geddes and Baker, 1967) tissues. Finally, the computed forward model and scalp topography for each EEG source were used to estimate the location of a single equivalent dipole model for each source (**Figure 1D**).

To co-register dipoles across subjects, warping parameters (**Figure 1E**) mapping each subject’s brain volume into Montreal Neurological Institute (MNI) space were estimated using the AFNI program *@auto_tlrc* (Cox, 1996). Then, parameters were applied to dipole locations, resulting in all subjects’ source dipoles having a common anatomical reference (see **Figure 1F**).

#### Clustering of EEG sources across subjects

The 1,521 sources with “near-dipolar” projections (having less than 15% residual variance between dipole projection and scalp topography; see Delorme et al., 2012; McLoughlin et al., 2013) and having coordinates within the boundaries of the template brain were grouped across subjects using EEGLAB’s k-means clustering framework (Onton et al., 2006). Sixteen source clusters (i.e., 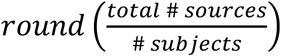; Rapela et al., 2012) were determined using the following features extracted from each source as contributing factors to the k-means algorithm: the MNI coordinates of equivalent current dipole locations, four principal components from normalized continuous EEG power spectra (chosen by each having accounted for more than 5% of the total variance in within normalized spectral power), and sixteen principal components (chosen to be equal to the selected number of clusters) from scalp topographical maps. To compensate for the low dimensionality inherent to dipole coordinates and increase their influence on the k-means solution, clustering dimensions corresponding to dipole coordinates were multiplied by a factor of five (cf. Gramann et al., 2010; Rapela et al., 2012; Piazza et al., 2016).

To keep clusters homogenous in terms of their contributing factors (i.e., dipole locations, power spectra, scalp topographies), sources located more than 2 standard deviations away from each cluster centroid in k-means measure space were removed from initial clusters. While this “robust” k-means process has been shown to return homogenous clusters (e.g., Ehinger et al., 2014; Knaepen et al., 2015; Lisi and Morimoto, 2015; Behmer and Fournier, 2016), it frequently returns clusters that are missing sources from one or more subjects. Therefore, to preserve sample size, comparable sources were added from subjects having “outlying” (i.e., > 2 std. dev.) sources in the k-means output (e.g., as in McLoughlin et al., 2013) to clusters of interest (i.e., those clusters associated with errors in the stimulus-response interval) based on having smallest Euclidean distances to centroids in the original clustering space.

Source time-series for each subject were forward projected (i.e., *X*_*channel*_ = *W*^−1^*u*_*source*_; see **Figure 1**) to scalp locations having the greatest topographical projections. For each time-series and within each subject, the electrical potential at each latency was standardized as ratio to baseline root mean square deviation within the concatenated -800 to -300 millisecond pre-stimulus intervals across all trials (cf. McLoughlin et al., 2013).

### Trial EEG measures

Of the 91 subjects producing both correct and error responses (two subjects made no errors), there were an average of 12.8 congruent (*SD* = 11.8) and 10.5 incongruent (*SD* = 9.7) error trials. Overall, response times (RTs) on error trials were shorter than on correct trials by about 58 milliseconds (*t*[36,350] = -14.63, *p* < .001 in linear mixed models [LMMs] accounting for multiple records within family and individual); RTs were also affected by stimulus incongruence (main effect of incongruent stimulus, *t*[36,332] = 32.46, *p* < .001; incongruent stimulus × error response interaction, *t*[36,335] = -7.76, *p* < .001). To make potentials for correct and error trials comparable, error trials were matched within subject to correct trials of the same stimulus type and nearest RT, giving a set of 4,196 trials (45.0% incongruent) for trial level analyses. This step was important to ensure that the relative duration and timing among stimulus- and response-related processes for a given error response trial and its comparison correct trial were similar; failure to adjust for variation in the duration of these processes may confound interpretation of associations between trial brain potentials and accuracy.

### Standard trial-average ERP and “regression-ERP” (rERP) measures

#### Standard trial-averaged ERPs

Stimulus- and response-locked brain potentials were separately computed by the “standard” ERP approach, which averages the potential across trials at each latency relative to stimulus or response, being largely sensitive to voltages of relatively consistent polarity at one or more latencies across trials. Standard ERPs for the four stimulus/response type combinations were averaged separately within subject. For comparability with other papers on action monitoring ERPs in youth (e.g., Pontifex et al., 2011; Torpey et al., 2012; Pontifex et al., 2013; Meyer et al., 2014; Anokhin and Golosheykin, 2015), subject waveforms were included if they possessed at least six artifact-free trials within a given stimulus/response type. As such, trial counts for the four trial types were as follows: correct congruent trials (*n* = 93 subjects, *M* = 249.5 [*SD* = 53.2]), correct incongruent (*n* = 93 subjects, *M* = 119.7 [*SD* = 27.9]), error congruent (*n* = 63 subjects, *M* = 17.1 [*SD* = 11.7]), error incongruent (*n* = 53 subjects, *M* = 15.6 [*SD* = 9.7]).

#### Overlap-corrected rERPs

When the time between stimulus and response varies across trials, standard ERPs “smear” overlapping stimulus- and response-locked brain potentials, such that some effect of potentials elicited by the stimulus are contained in the response-locked ERP and vice versa. Thus, “smearing” unsatisfyingly interjects uncertainty into interpretation of stimulus-versus response-related waveform features (Woldorff, 1993; Salisbury et al., 2001), making difficult identification of potentials unique to *either* stimulus *or* response. An alternative to standard ERP averaging is to decompose dissociable stimulus- and response-locked brain potentials via the “regression-ERP” (or rERP) framework (see methods papers by Smith, 2011; Burns et al., 2013; Smith and Kutas, 2015a, b; Ehinger and Dimigen, 2018), an overlap-correction application similar to the deconvolution method used in event-related fMRI research (Hinrichs et al., 2000) that enables insight into what separable stimulus- and response-locked processes contribute to standard ERPs in which these two processes are confounded.

In simplest terms, the rERP can be described as a vector of *β* coefficients from a series of regressions conducted separately at each latency within an epoch relative to an event. As detailed by Smith and Kutas (2015a, b) the standard ERP is a special case of the rERP where in the model

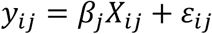

the observed EEG data *y* on trial *i* at latency *j* relative to a task event (e.g., stimulus onset) is the result of the “true” ERP *β*_*j*_ summed with un-modeled zero-mean “noise” ε at that latency; here, the trial design column matrix *X* consist entirely of ones with length equal to the number of trials. Equivalently, rather conducting separate regressions sequentially at each latency *j* within the event-locked epoch of length *J*, the many regressions can be combined into a single model to estimate all *β*_1,2,…,*J*_ simultaneously (software for applying the rERP technique outlined by Smith and Kutas, 2015a, b can be found at: *vorpus.org/rERP, sccn.ucsd.edu/wiki/EEGLAB/RERP*, and `unfoldtoolbox.org`). Specifically, the above column matrix *X* may be expanded with zeros to generate an *L*-by-*J* matrix *STIM* that accounts for all latencies *L* in the continuous EEG time-series (see **Figure 2A**). Here, 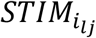 takes the value of 1 when there is intersection between the conditions of *l* and *j* (e.g., 300 milliseconds post-stimulus in the continuous EEG on trial *i and* 300 milliseconds post-stimulus in the event-related epoch), and 0 otherwise.

**Figure 2.**
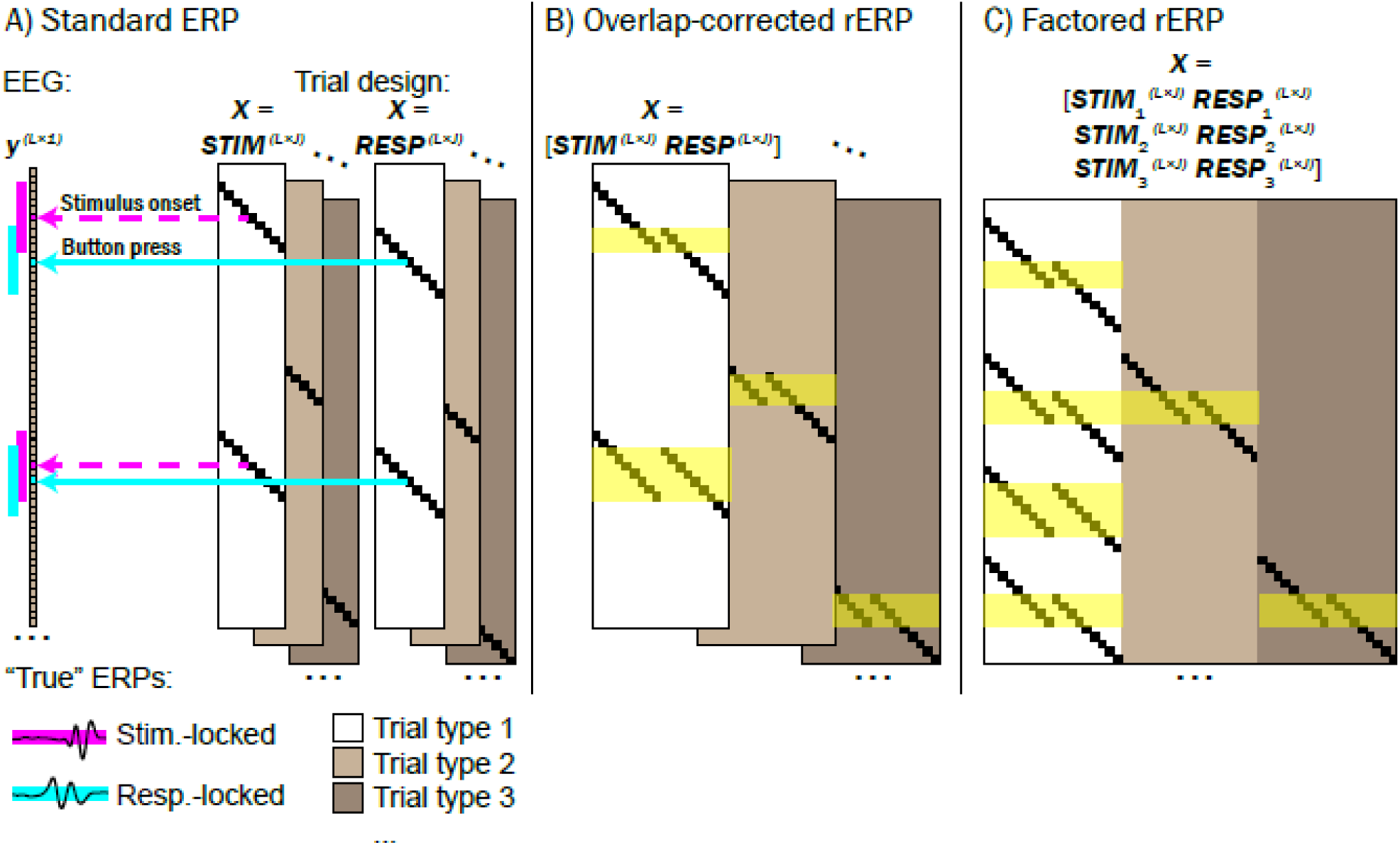
Comparison between standard trial-averaged ERP, overlap-corrected regression-ERP (rERP), and overlap-corrected factored rERP, using regression design. A) Standard ERPs time-locked to stimulus onset and button response events may be calculated separately by using separate *STIM* and *RESP* matrices substituted for the trial design matrix *X*, separately across different trial types (e.g., congruent correct, incongruent correct, congruent error, etc.). Here, each row of *X* corresponds to each latency *L* in the continuous EEG recording *y*, and each column of *X* corresponds to each latency within the chosen epoch for which the ERP is to be calculated (e.g., 2 seconds before and after each stimulus or response event) of length *J*. An element of *X* takes the value of 1 (small black squares) if it corresponds to a latency within the event-related epoch on a given trial number of a given trial type, and 0 otherwise. As trial RTs vary, so do overlapping non-zero elements of *STIM* and *RESP*. Note that the standard ERP does not account for temporal overlap among stimulus and response epochs (see magenta and cyan demarcations alongside the EEG recording *y*), and therefore stimulus- and response-locked standard ERPs are temporally confounded. B) Overlap-correction among stimulus and response processes is achieved by horizontal concatenation of *STIM* and *RESP* into a single matrix *X* for each trial type. Here, periods within the continuous EEG recording where stimulus- and response-locked epochs overlap are highlighted in yellow. C) Concatenating task design matrices from multiple trial types may be used to explore processes that are unique to a given trial type. Using trial type 1 (white block of *X*) as the reference type in a treatment coding framework, the degree to which waveforms deviate from trial type 1 as a function of trial type 2 (light gray) or trial type 3 (dark gray) may be explored. For example, in our study we used congruent correct trials as the reference type (white block) and explored how rERPs deviated as a function of incongruent flanker stimuli (light gray) or errors (dark gray).

Overlap in stimulus- and response-locked brain potentials may then be modeled by horizontal concatenation of the above design matrix *STIM* with a similar design matrix *RESP* that is conditioned to button response events (see **Figure 2B**). For example, in the equation

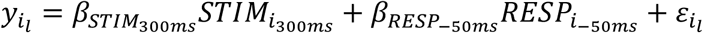

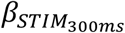 and 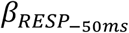 are “true” potentials co-occurring 300 milliseconds after stimulus presentation and 50 milliseconds before the button response (respectively) on trial *i*.

Mirroring the approach taken for standard ERPs, overlap-corrected rERPs were computed for the four stimulus/response combinations. Overlap-corrected rERPs enabled straightforward interpretation of separate stimulus-versus response-related processes, such that they provide an estimate of what overlapping stimulus- and response-locked brain potentials separately contribute to standard trial-averaged ERPs.

#### Overlap-corrected, factored rERPs

Taking our rERP investigation further, we used a treatment coding approach (cf. Smith and Kutas, 2015a, b) to highlight brain potentials associated with stimulus and response “factors.” This strategy is similar to calculating “difference waves” (cf. Luck, 2005), whereby the standard waveform of one or more stimulus/response combinations (e.g., correct trial) may be subtracted from another (e.g., error trial), although the key advantage to factored rERPs is that they permit overlap-correction. Subtraction of scalp-recorded channel waveforms from different task conditions may distort true brain activity because the spatiotemporal sum of active sources projecting to a scalp channel at one task moment may differ from that during other moments (e.g., see discussion in Burwell et al., 2016, p. 1003). But, we believe the approach of using ICA-unmixed component processes circumvents these concerns because contrasts are made on EEG *source activities* themselves.

To isolate stimulus- and response-locked rERP task factors (see **Figure 2C**), we extended the above rERP model to:

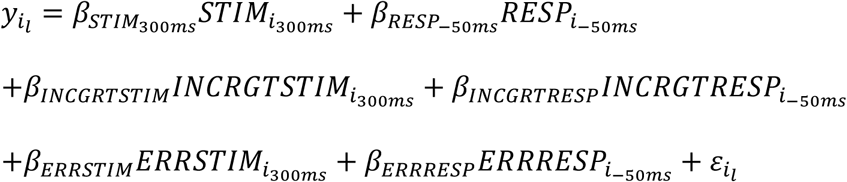

As before, *β*_*STIM*_ and *β*_*RESP*_ reflect respective stimulus- and response-locked rERPs, but here they are fixed to a chosen reference, which we coded to be congruent correct trials. Then, in the full system of equations, *INCGRTSTIM*_*i*·_ / *INCGRTRESP*_*i*·_ and *ERRSTIM*_*i*·_ / *ERRRESP*_*i*·_ are coded 0 (for congruent trials and correct responses, respectively) and 1 (for incongruent trials and error responses, respectively). Critically, the resulting waveforms comprised by *β*_*INCGRTSTIM,ERRSTIM*_ and *β*_*INCGRTRESP,ERRRESP*_ do not reflect trial waveforms, but rather the expected time-varying *deviations* from waveforms *β*_*STIM*_ and *β*_*RESP*_ as a function of task stimulus and response type. We derived three stimulus-locked and three response-locked waveforms for each subject to reflect: 1) the expected brain potential on correctly-performed congruent stimulus trials, 2) deviation when flanking stimuli are incongruent with the target stimulus, and 3) deviation associated with erroneous responses. We explored an incongruence-by-error interaction, but determined its impact nonsignificant; thus, here we only estimated main effects of stimulus incongruence and response errors.

### Experimental Design and Statistical Analyses

#### Within-subject prediction of errors using EEG, ERP, and rERP

Voltages from single-trial EEG, standard trial-averaged ERPs, and overlap-corrected rERPs were used as predictors of response errors (separately for congruent and incongruent trial types), in the following logistic regression linear mixed model (LMM; Pinheiro and Bates, 2000): *ERROR*_*ijk*_ = *B*_*INT*_ + *B*_*ERROR*_*VOLTAGE*_*ijk*_ + α_*jk*_ + α_*k*_ + ε_*ijk*_. Here, *ERROR*_*ijk*_ and *VOLTAGE*_*ijk*_ respectively reflect response errors (1 = error, 0 = correct) and voltages (from baseline-subtracted single-trial EEG, trial-averaged ERP, or overlap-corrected rERP) on trial type *i* for participant *j* of family *k, B*_*INT*_ is the model intercept, ε_*ijk*_ is random noise, and random intercept terms α_*jk*_ and α_*k*_ account for multiple records per subject (i.e., trials) and multiple records per family (i.e., twins), respectively. For single-trial EEG, logistic regressions were conducted at each latency relative to stimulus or response events; for trial-averaged ERPs and overlap-corrected rERPs, logistic regressions were conducted for mean potentials extracted from windows centered around peaks [±50 milliseconds] identified as relevant for errors in single-trial analyses. The reported fixed-effect coefficient *B*_*ERROR*_ reflects the expected change in probability (log odds) of an error with one standard deviation positive-going change in EEG, ERP, or rERP. Odds ratios (*OR*s, presented in tables and calculated by exp[*B*_*ERROR*_]) reflect the proportion increase in the odds of an error, adjusted for nuisance covariates (Agresti, 2013). P-values for logistic regressions were determined by calculating the z-values for each *B*_*ERROR*_ coefficient (i.e., 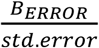) and referring it to the standard normal quantile function. For significance testing of within-subject analyses (logistic regression prediction of errors) we used a false discovery rate-corrected (FDR, Benjamini and Hochberg, 1995) alpha of .01; between-subjects analyses (prediction of individual differences in error rate, ADHD analyses) adopted a more sensitive FDR-corrected alpha of .05, which were flagged in tables using asterisks. Although not shown in the model and not reported, single-trial or trial-averaged RT, subjects’ age, and gender were included as fixed-effect nuisance covariates.

Source clusters whose ERP waveforms within the SRI were found to significantly predict errors across both congruent and incongruent trial types were further examined next to lateralized readiness potentials (LRPs), which are thought to reflect activation of motor preparation and execution processes and indicate the responding hand by greater contralateral than ipsilateral hemisphere negativity (Smulders and Miller, 2011). This enabled comparing the temporal sequencing between brain processes generally related to errors and processes reflecting motor activation of the correct or erroneous button response; whichever began first may be considered be more central to error commissions. Like above, LMM logistic regressions predicting errors were calculated at each latency, but instead of analyzing separately congruent and incongruent trials, analyses were conducted separately within trials grouped by the correct-response hand (including a stimulus congruency interaction term). LRPs were calculated in the typical fashion (de Jong et al., 1988; Gratton et al., 1988) by subtracting the ipsilateral scalp waveform (e.g., CP4 for right-hand responses) from that of the contralateral scalp waveform (CP3 for right-hand responses), but unlike most prior investigations studying LRPs, scalp voltages were summed projections only from independent component source clusters in or near left and right sensorimotor cortex (thereby spatially filtering out influences from other sources). We compared the relative timings of error-related effects (*p*_FDR_ < .01) for non-LRP and LRP voltages to understand which manifested first.

#### Between-subject prediction of error rates using factored rERPs

We next sought to understand whether brain potentials reflected in factored rERP waveforms predicted individual

differences in task error rates (i.e., 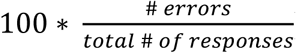). Error rates for the overall sample were positively skewed, so to approximate a normal distribution to be used in linear regressions, error rates *x* were transformed by log(*x* + 1). Next, at each latency within the factored rERPs, the following LMM was estimated: *PCTERR*_*ijk*_ = *B*_*INT*_ + *B*_*PCTERR*_*VOLTAGE*_*jk*_ + α_*jk*_ + α_*k*_ + ε_*jk*_. In this equation, *PCTERR*_*jk*_ and *VOLTAGE*_*jk*_ reflect error rate and baseline-subtracted potential at a given latency relative to either stimulus or response for subject *j* of family *k*.

#### Do amplitude differences in rERPs mediate the association between ADHD symptom count and heightened task error rates?

Possible brain mechanism(s) responsible for the association between ADHD and high task error rates remain unknown. Given an association between brain potentials and error rates, brain potentials may plausibly be tested as indicators of the relationship among ADHD symptom count and error rates. Specifically, in the mediation framework (for review, see Shrout and Bolger, 2002): *Does the predictor (ADHD symptom count) exert its statistical effect on the outcome (error rate) by way of a mediator (brain potential measure), such that when both predictor and mediator are included as covariates, the effect of the predictor variable is significantly reduced?*

We first examined whether ADHD symptom counts, stimulus congruence (incongruent vs. congruent), and congruence × ADHD symptom count interaction significantly (*p* < .05) predicted task error rates in LMMs (including age and gender as fixed-effect covariates and a random intercept to account for multiple records per family). In mediation framework, the main effect of ADHD on error rates may be thought of as the “total” effect, as illustrated by path *c* in **Figure 3A**. However, the “total” effect of ADHD on error rate may be partitioned into “direct” (path *c’*, **Figure 3B**) and “indirect” effects (paths *a* and *b*). The direct effect *c’* reflects the reduction from the total effect to the indirect effect, which is mathematically equivalent to the product of *a* and *b*.

**Figure 3.**
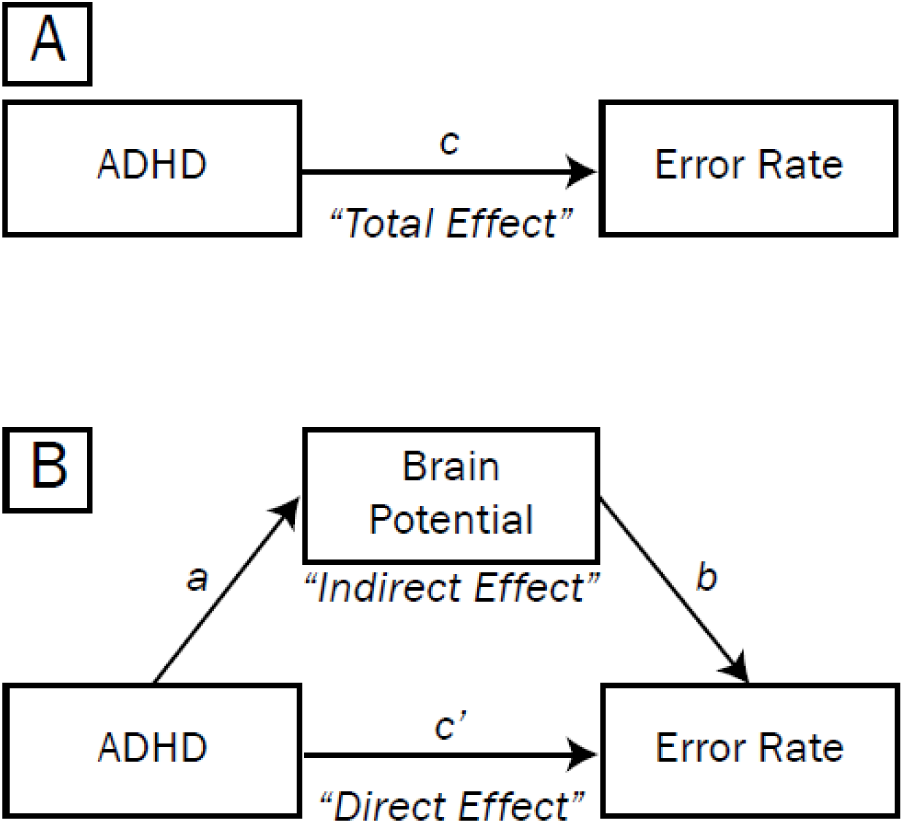
Mediation model testing whether brain potentials account for the association between ADHD symptom count and task error rates. The main effect of ADHD symptoms on error rates may be thought of as the “total effect”, as illustrated by path *c* (A). However, the “total” effect of ADHD on accuracy may be partitioned into “direct” (path *c’*, B) and “indirect” effects (paths *a* and *b*). The direct effect *c’* reflects the reduction from the total effect to the indirect effect, which is mathematically equivalent to the product of *a* and *b*.

In the mediation model, a brain potential is said to have partially mediated the relationship between ADHD symptom counts and error rates if the shrinkage in the value of *c* to *c’* is significant; or equivalently, if the size of the indirect effect *ab* is significantly different than zero. To quantify paths *a* and *b*, for each subject we extracted mean potentials(s) within time segments of the factored rERP waveforms that: (1) were strongly predictive of overall task error rate (requiring *p*_FDR_ < .05 for a duration of at least 50 consecutive milliseconds in the rERP waveform), and (2) overlapped temporally with EEG/ERP time windows determined by above analyses to robustly predict response errors. These measures were separately tested as mediators (cf. “Brain Potential” in **Figure 3B**) in the association between ADHD symptoms and task error rates. Paths *a, b, c*, and *c’* were presented as *t*-statistics (i.e., 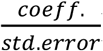), and denominator degrees of freedom were calculated using Kenward-Roger approximation (Kuznetsova et al., 2016).

To estimate whether mediation was significant, we used a bootstrapping approach used by Burwell et al. (2014) whereby 1,000 indirect effects (*ab*, equivalent to the change in *c* to *c’*) were simulated from the observed data with replacement (Shrout and Bolger, 2002), keeping the proportion of matched- to unmatched-twin-pairs constant. Ninety-five percent confidence intervals were estimated from this distribution of simulated *ab* effects to evaluate statistical significance at the .05 level; significant mediation is said to have occurred when the confidence interval did not include 0.

## Results

### Predicting errors within-subject using single-trial source potentials

In single-trial analyses, correct and error RTs did not differ (*t*[4,103] = -.72, *p* = .470, LMM linear regression), nor was the interaction with stimulus incongruency significant (*t*[4,103] = -.34, *p* = .731, LMM linear regression), confirming that the matching procedure for single-trial analyses was successful and that the relative timing among stimulus- and response-related processes for a given pair of correct and error trials within subject was similar.

In **Figure 4**, we plotted grand mean waveforms for each trial type derived from fifteen source clusters (one cluster not shown because it corresponded to non-brain eyeblink artifacts) as well as mean dipole locations. Horizontal bars are plotted above waveforms to highlight periods when the source electrical potential derived from individual trials significantly predicted an error response within congruent or incongruent stimulus trials (*p*_FDR_ < .01 for a period of 50 milliseconds or longer, LMM logistic regression). Several error-related effects were detected, such as greater post-error negativity (in medial frontal clusters, approximately 0 to 100 milliseconds post-response) and greater post-error positivity (mainly in clusters focused in posterior medial frontal and parieto-occipital regions, approximately 100 to 600 milliseconds post-response) in response-locked waveforms. However, the only source cluster for which we observed significant and consistent error associations across congruent and incongruent trials within the stimulus-response interval (i.e., the mean SRI between the dashed and solid vertical lines) was a cluster located primarily in posterior medial frontal cortex (pMFC, see blue arrows in **Figure 4**). The mean scalp projection and subject dipoles for pMFC are plotted in **Figure 5**, which were primarily focused in bilateral mid-cingulate cortex (39%) and supplementary motor area (21%) of the common template atlas (Tzourio-Mazoyer et al., 2002). In **Figure 6**, source contributions from this cluster (blue envelopes) to grand mean potentials across all channels (the mixture of all sources’ and clusters’ contributions, outer gray traces) are depicted, and accounted for approximately 5% to 14% of the variance of voltages reflected in scalp channels.

**Figure 4.**
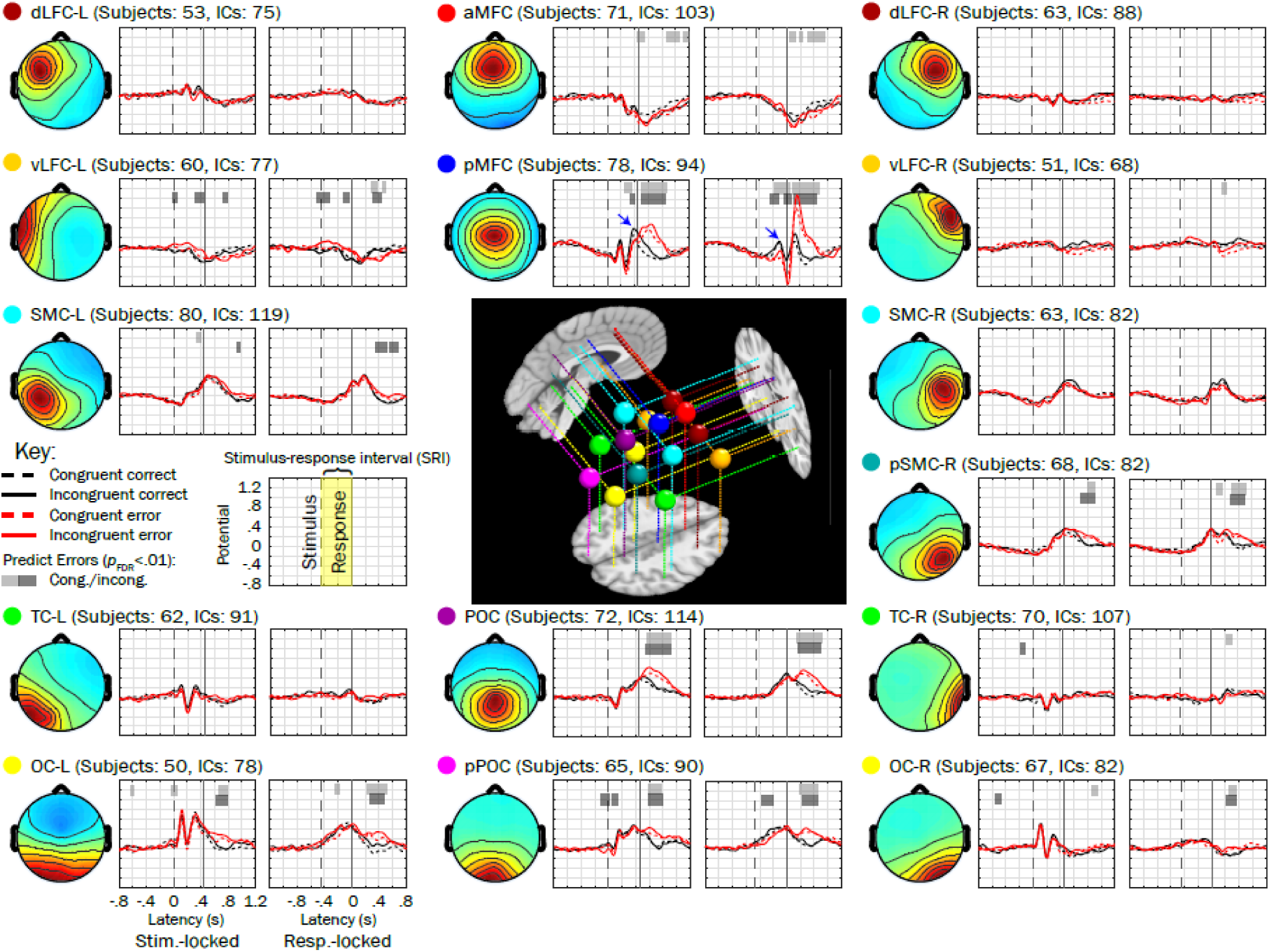
Fifteen source clusters, grand mean trial waveforms, and voltage associations with errors. Grand mean waveforms from error (red traces) and correct (black traces) performance trials (matched within stimulus congruency types [congruent = dashed, incongruent = solid] by response time) are plotted next to mean scalp topographies for each of 15 source clusters (one of the clusters not shown because it primarily reflected ocular activity) examined in the present study. Horizontal bars plotted above waveforms indicate regions where the mixed model logistic regressions significantly predicted the occurrence of an error (having *p*_FDR_ < .01 for more than 50 consecutive milliseconds); light shaded bars correspond to congruent trials whereas dark bars correspond to incongruent trials. We looked for significant associations within the mean stimulus-response interval (SRI, labeled in the key). Mean dipoles for each of these clusters are plotted in the center on representative sagittal, axial, and coronal slices from the template brain. Abbreviations: dLFC = dorsolateral frontal cortex; MFC = medial frontal cortex; vLFC = ventrolateral frontal cortex; SMC = sensorimotor cortex; TC = temporal cortex; POC = parietooccipital cortex; OC = occipital cortex; the prefixes “a” and “p” refer to “anterior” and “posterior” (respectively) and suffixes “-L” and “-R” correspond to left and right hemispheres.

**Figure 5.**
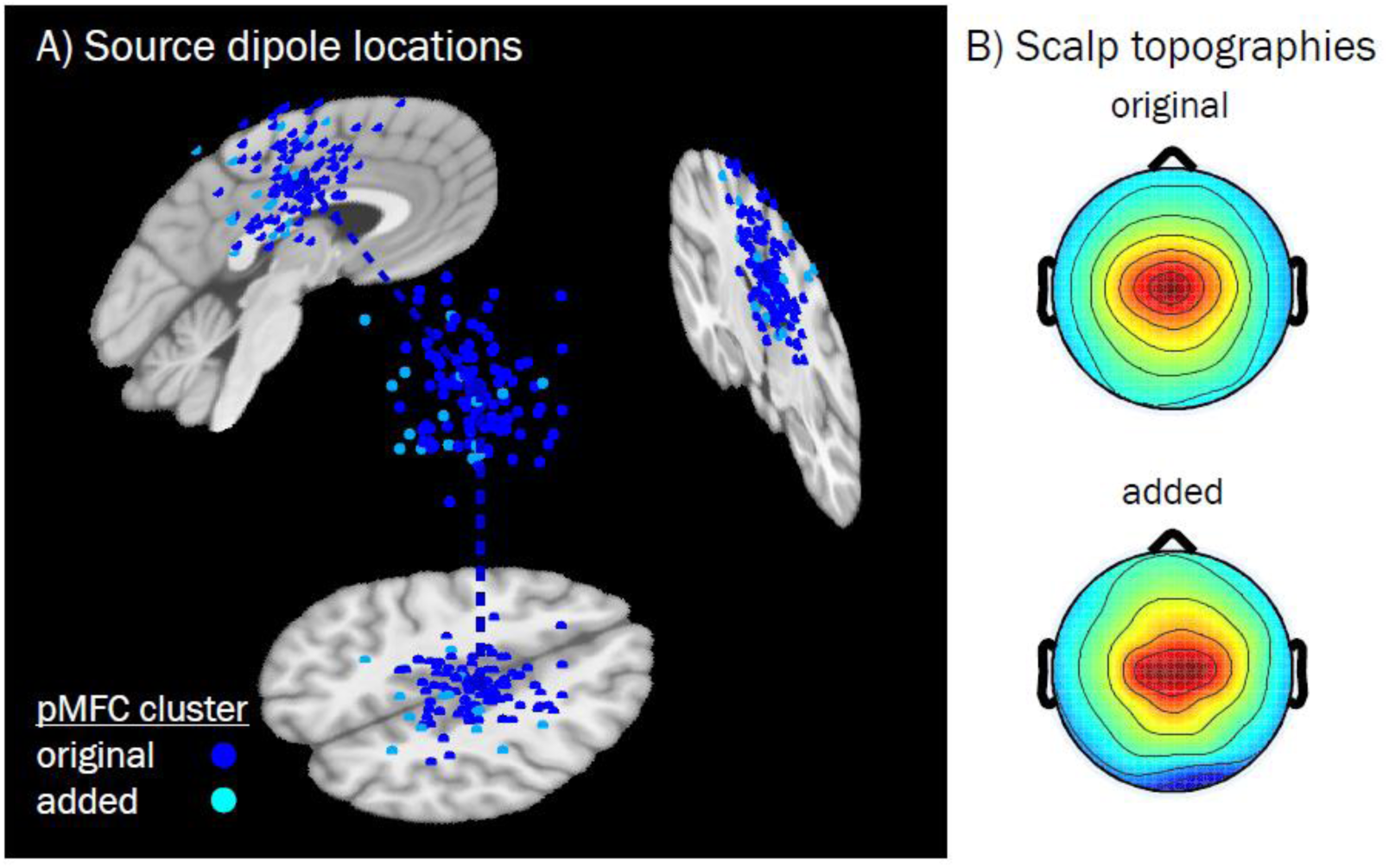
Equivalent current dipoles and mean scalp topographies for the pMFC source cluster derived by the k-means algorithm and post-hoc additions. A) Subject dipoles and their projections onto MNI template space for pMFC are smaller in size than the cluster centroid (located in right mid-cingulate cortex, MNI coordinates: 4, -12, 44), which is larger. Blue subject dipoles depict sources that were included in the original k-means clustering output (*N* = 94, across 78 subjects); cyan dipoles correspond to sources that were added (for subjects with no dipole in the original cluster, *N* = 16) subsequently based on nearest distance in clustering measure space. B) Mean scalp topography for sources that were originally included in the k-means obtained pMFC cluster (above), and the topography for those sources which were subsequently added (below). Added subjects did not differ from the original pMFC-clustered subjects in terms of demographics (female vs. male: *t*[22] = -1.10, *p* = .284; age: *t*[21] = -.72, *p* = .477, Welch’s two-sample t-test) or study variables of interest (error rate [log-transformed]: *t*[78] = 1.00, *p* = .319; ADHD symptoms [log-transformed]: *t*[78] = -1.48, *p* = .142, LMM linear regression). Note that while some dipole coordinates appear as being located outside of the brain for a given sagittal, coronal, or axial MRI slice (slices chosen based on the centroid coordinates), all dipoles were contained within the boundaries of the template brain as part of their inclusion criteria.

**Figure 6.**
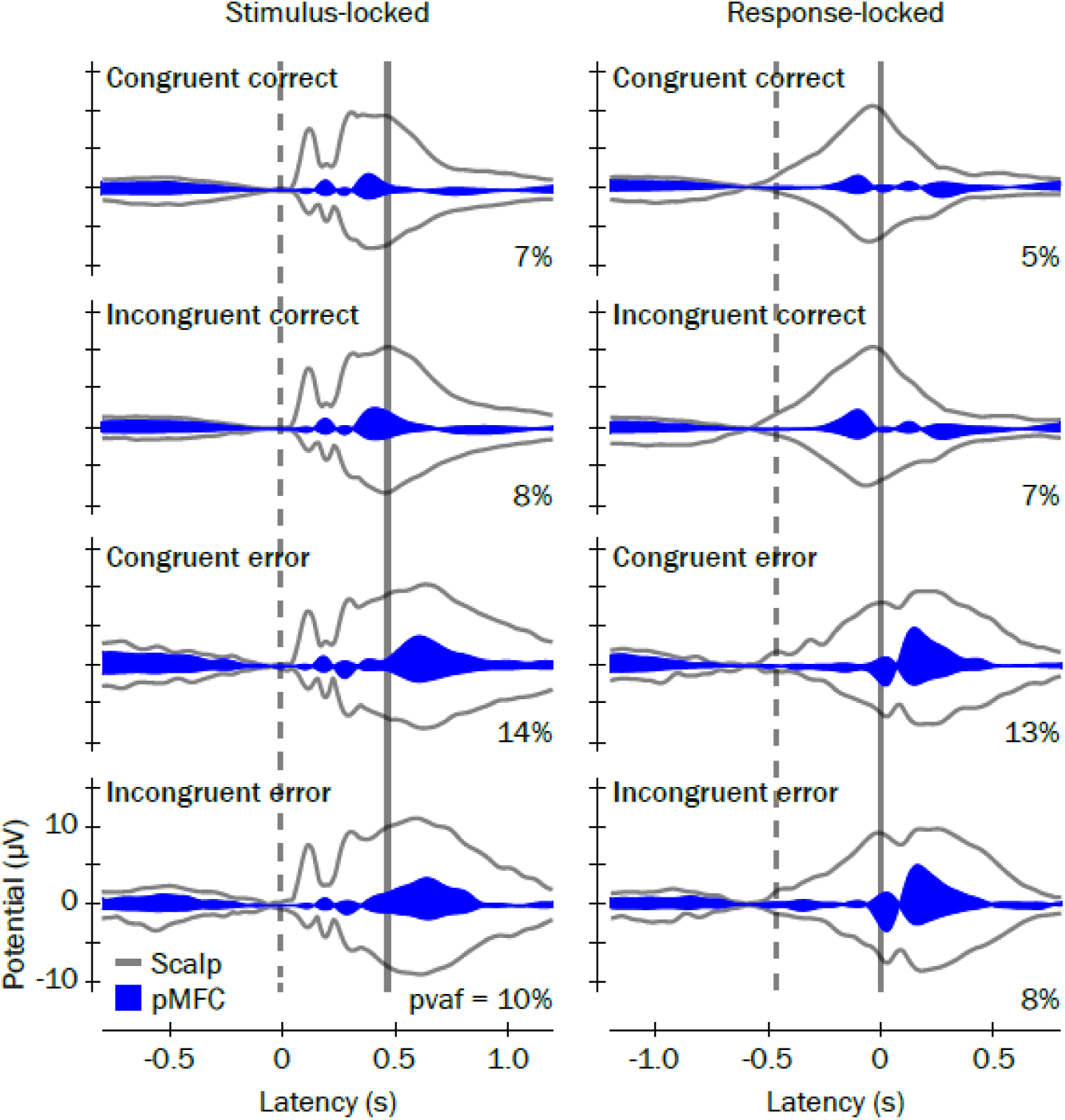
ERP envelopes and contributions of the pMFC cluster. ERP envelopes (outer gray traces) show the maximum and minimum scalp channel voltages for each latency in the grand mean ERP waveforms, time-locked to stimulus presentations (left) and button responses (right). Grand mean latencies for stimulus presentations and button responses are denoted with vertical dashed and solid lines, respectively. Contributions of the pMFC source cluster to the grand mean ERPs are shown as blue envelopes and the percent variance accounted for 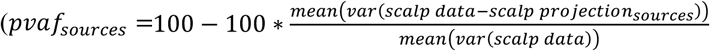) by pMFC to each ERP envelope are reported.

We identified two positive-going peaks within pMFC-derived source waveforms for which the voltage within the SRI was more positive in correct performance trials than in error trials. These peaks occurred at approximately 390 milliseconds after stimulus presentation in stimulus-locked waveforms and 110 milliseconds before the response in response-locked waveforms and are labeled in **Figure 7A**. Because of the frontocentral midline scalp distribution of pMFC and the timings of these positive-going peaks relative to correct trial stimulus and response events, we termed these stimulus- and response-locked peaks as *Frontocentral P3* and *Pre-Movement Positivity (PMP)*, respectively.

**Figure 7.**
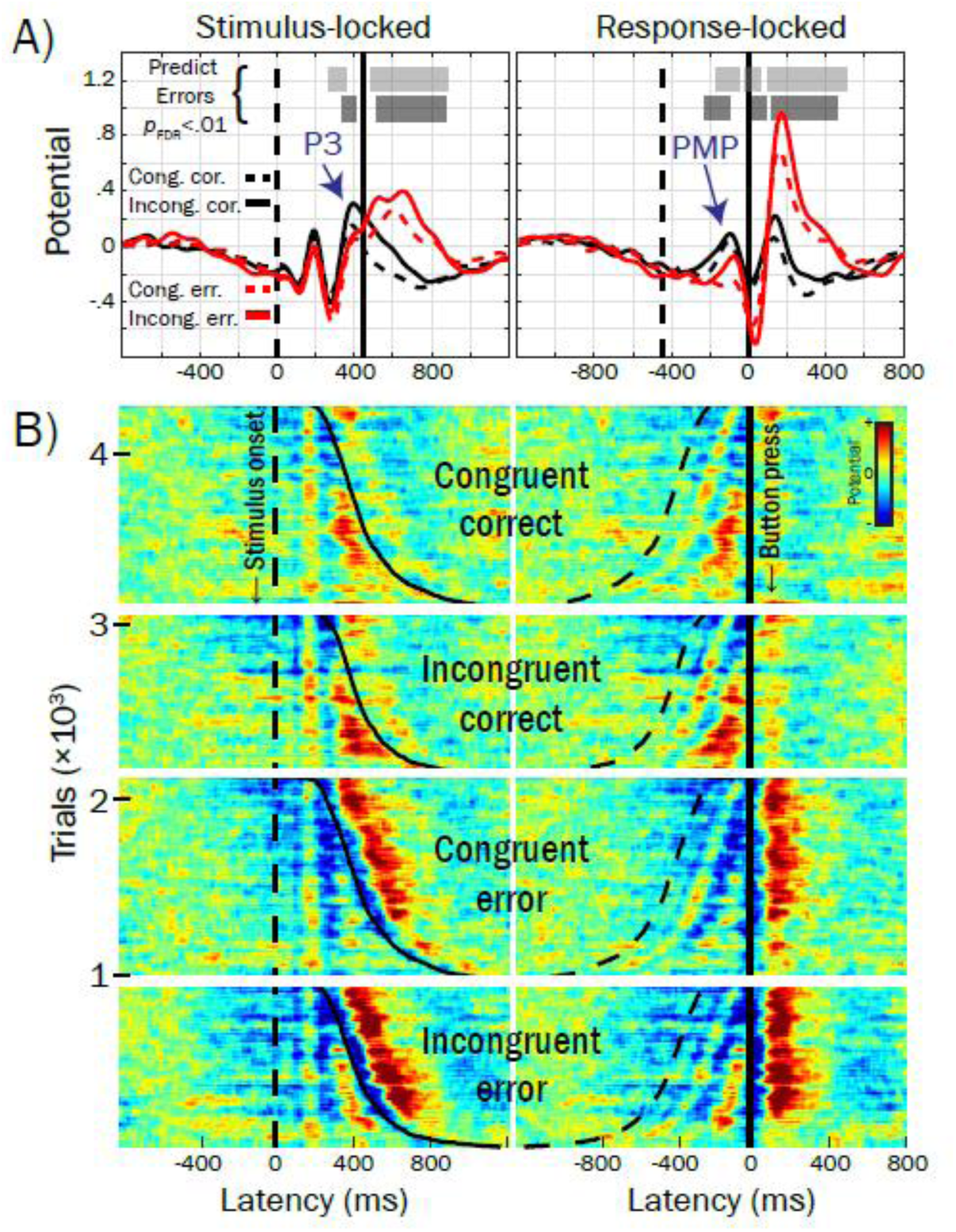
Grand mean waveforms from response-time-matched error and correct trials for the pMFC cluster activities and single-trial ERP-images. A) Voltage peaks within the mean stimulus-response interval (SRI, i.e., between the vertical dashed [stimulus] and solid [response] lines) that were associated with errors in single-trial logistic regressions are labeled: Frontocentral P3 and Pre-Movement Positivity (PMP). B) ERP-images for response-time-matched error and correct trials, across subjects. Stimulus- and response-locked ERP-images are labeled, reflecting time-varying voltage (referenced and scaled to baseline, see text) for each trial stacked and sorted by stimulus congruency (congruent, incongruent), response accuracy (error, correct), and response time (dashed line = stimulus-onset, solid line = button-press); red colors reflect positive voltage deflections and blues reflect negative voltage deflections (root mean square normalized within subject) forward-projected to channel Cz. Trials in ERP-images are grouped by congruent correct (*N* = 1,193), incongruent correct (*N* = 965), congruent error (*N* = 1,193) and incongruent error (*N* = 965). For example, the first row in the first group of trials corresponds to a correctly-performed congruent trial drawn from the subject with the fastest RT across all subjects; the last row in the first group of trials corresponds to a correctly-performed congruent trial from the subject with the slowest RT across all subjects. For ease of visualization, ERP images were smoothed vertically using a sliding boxcar window of 1% width the overall number of trials (*N* = 4,196).

We also examined error-related Frontocentral P3 and PMP temporal relations to lateralized readiness potentials (LRPs) thought to reflect central motor activation of the button press, enabling insight into whether error-related pMFC activations preceded brain activations reflecting the specific motor response to be carried out (i.e., left or right finger presses). In **Figure 8A**, grand mean waveforms averaged within correct (dashed traces) and error (solid traces) trials, separately for trials where the correct response required either left (green) or right (blue) button presses, are plotted for the pMFC cluster, as well as left (SMC-L) and right (SMC-R) clusters reflecting putative sensorimotor sources that were projected to scalp sites where LRPs are typically recorded. In **Figure 8B**, LRPs peaking approximately -75 milliseconds in the response-locked ERP encode the overt button response and are labeled with blue arrows: on correct trials, the LRP is negative-going contralateral to the finger press (reflecting activation of the correct response), whereas on error trials, the LRP is a positive-going peak in the contralateral hemisphere (reflecting incorrect motor activation). PMP and LRP voltages predicted errors at overlapping latencies, although PMP effects began earlier (−180 vs. -148 milliseconds in response-locked waveforms), suggesting that error-related pMFC processes accompany and may begin before erroneous motor activations reflected in SMC.

**Figure 8.**
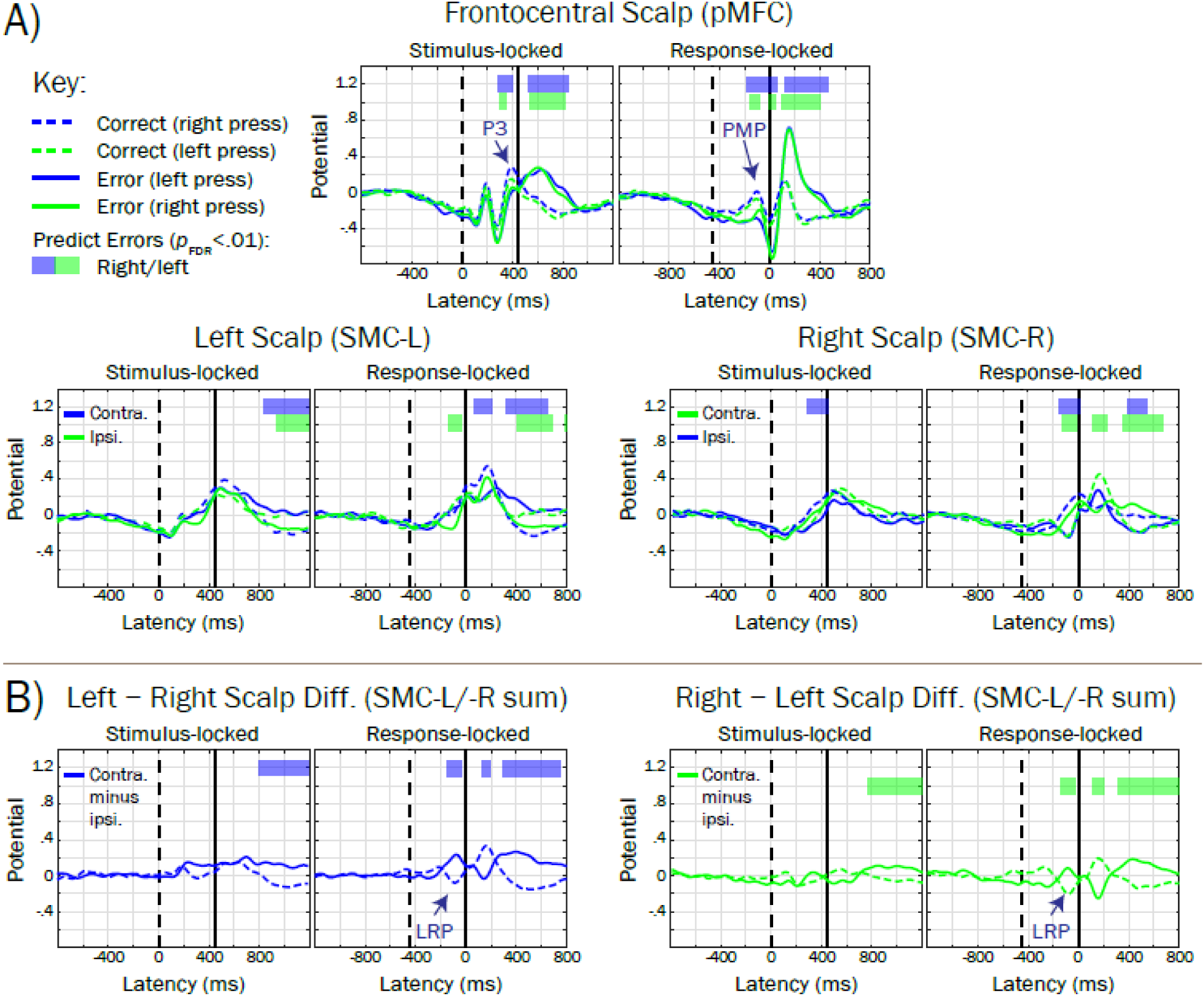
Grand mean trial waveforms reflecting posterior medial frontal cortex (pMFC) and lateralized motor-related potentials, and their association with errors. A) Grand mean waveforms for the pMFC source cluster and left (SMC-L, projected to site CP3) and right (SMC-R, CP4) clusters putatively reflecting sensorimotor cortical sources are averaged within trial accuracy (dashed trace = correct; solid trace = error) and the correct-response hand mapping (blue trace = right hand; green trace = left hand). Horizontal bars denote regions of the waveform that significantly predict errors (blue = correct-response right hand; green = correct-response left hand), accounting for a stimulus congruency interaction term. For comparability, trials and subjects contributing to SMC-L and SMC-R waveforms were identical to that of the pMFC cluster (unmatched subjects were included based on Euclidean distance in clustering measure space, see Methods and Materials). B) Lateralized readiness potentials (LRPs, labeled with navy arrows) were revealed by summing the scalp projections for SMC-L and SMC-R clusters, and subtracting the ipsilateral from contralateral waveform, consistent with guidelines reviewed by Smulders and Miller (2011). Note that the polarity of the LRP peak (approximately -75 milliseconds in response-locked waveforms) encodes the response hand mapping, such that it is negative-going for contralateral responses and positive-going for ipsilateral responses.

In **Figure 7B**, stimulus- and response-locked potentials across trials from the pMFC source cluster (across subjects) are visualized by ERP-image. Here, each row (horizontal colored line) corresponds to potential in a single trial plotted over time relative to stimulus and/or response events; red and blue hues indicate positive and negative deflections, respectively. Frontocentral P3 and PMP peaks in **Figure 7A** appear in warm hues (reds, yellows) that are consistent across rows at the same latencies in ERP-images in **Figure 7B**. Importantly, ERP-images illustrate the fact that button presses (solid traces) frequently occurred within close temporal succession of the presentation of the stimulus (dashed traces), resulting in overlap and possible confounding among Frontocentral P3 and PMP.

### Predicting errors within-subject using trial-summated ERP and rERPs

Typically, single-trial EEG traces are sufficiently variable that trial-to-trial brain potentials are difficult to discern and thus are not readily usable in between-subject analyses. Thus, we followed up single-trial analyses with analyses conducted on standard trial-average ERPs and overlap-corrected rERPs, which do afford these capabilities. In **Figure 9**, waveforms are plotted showing (**A**) standard ERP trial averages and (**B**) overlap-corrected rERPs. Despite having comparable shapes, standard ERPs and overlap-corrected rERPs are not identical, as illustrated by horizontal bars reflecting latencies at which the amplitude of ERP/rERP waveforms deviated (*p*_FDR_ < .01) from the mean potential within the pre-stimulus baseline. Further, subtraction of rERP estimates from trial data adequately accounted for evoked potentials (see **Figure 10A-D**), whereas subtraction of standard ERPs from trial data does not and may introduce artifact (compare **Figure 10D-E**). **Figure 9C** depicts the factored rERP waveforms for the expected ERP on congruent correct trials (black dashed trace), the event-related deviation when flanker stimuli are incongruent with the target stimulus (black solid trace), and the event-related deviation associated with erroneous responses (red trace). Factored rERPs highlight processes specific to stimulus type and response accuracy factors, which are confounded in **Figure 9A-B**.

**Figure 9.**
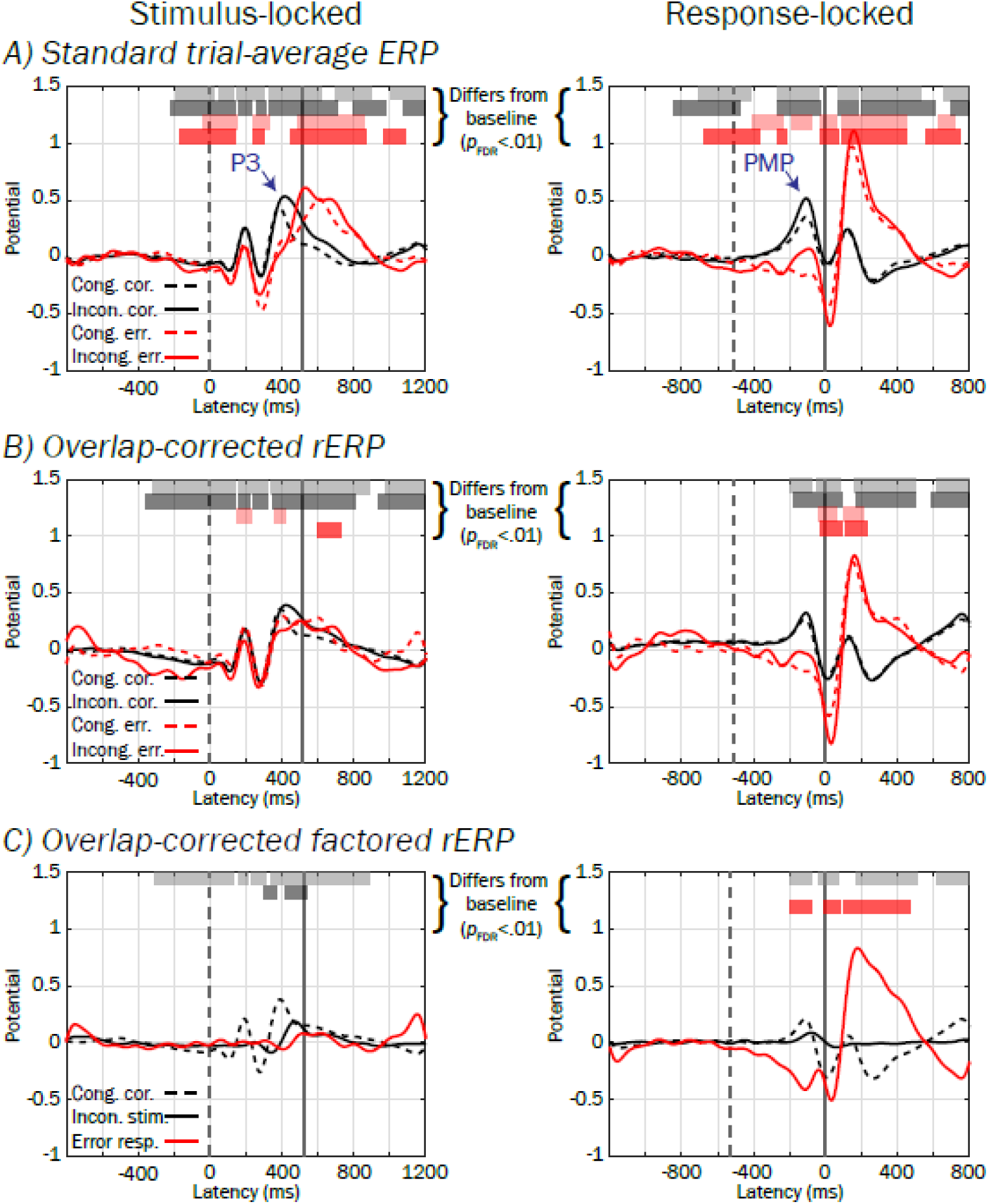
Standard ERPs, overlap-corrected rERPs, and overlap-corrected factored rERPs from the pMFC cluster. A) Standard trial-averaged stimulus-(left) and response-locked (right) ERPs, time-varying voltage of relatively consistent polarity at one or more latencies across trials (scaled to a pre-stimulus baseline period, see text) for each stimulus (congruent stimuli = dashed traces; incongruent = solid traces) and response type (correct = black; error = red). B) Overlap-corrected regression ERPs (rERPs, described in text) can be similarly interpreted to standard ERPs, although unlike standard ERPs that confound stimulus- and response-locked potentials occurring in the same time window, rERPs provide an estimate of what stimulus- and response-locked ERP processes separately contribute via summation to the time-overlapped standard ERP. C) Overlap-corrected, factored rERPs show the potential on congruent correct-performance trials (black dashed trace), 2) deviation when flanking stimuli are incongruent with the target stimulus (black solid trace), and 3) deviation associated with erroneous responses (red trace). The mean value within a pre-stimulus baseline interval was subtracted from all waveforms. Vertical dashed and solid lines denote mean stimulus and response latencies respectively.

**Figure 10.**
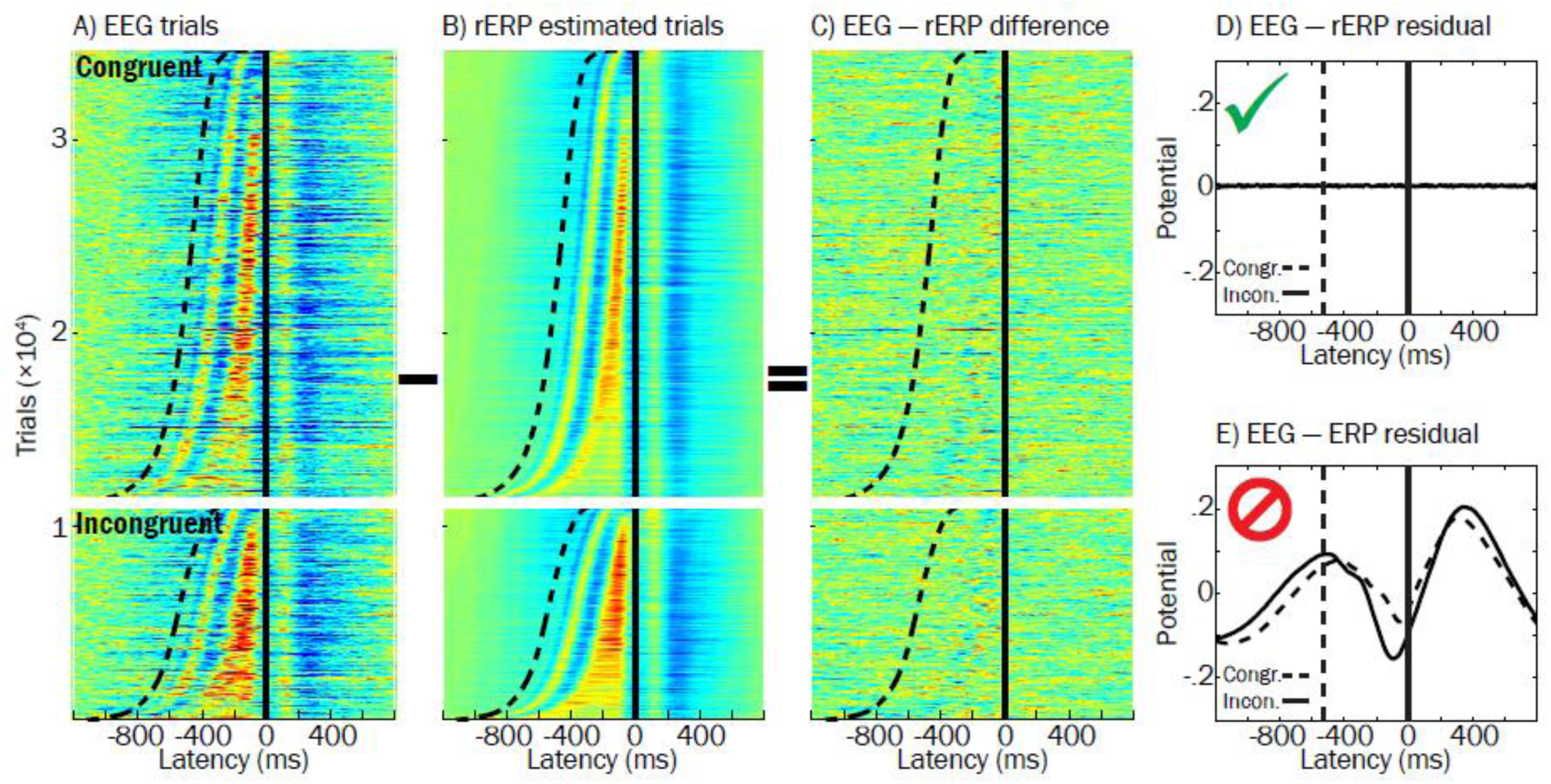
Effectiveness of using regression ERPs (rERPs) to account separately for stimulus and response processes that overlap in trial EEG. A) The “raw” response-locked ERP image for all congruent and incongruent hit trials is plotted, sorted by reaction time. Sigmoidal dashed line and vertical solid line reflect stimulus and response latencies, respectively. B) Pseudo-data for rERP estimated trials was constructed by: 1) substituting each subject’s response-locked rERP waveform (aligned to the button-press) in place of his or her trial data, and then 2) superimposing that subject’s stimulus-locked rERP, shifted backwards in time with respect to stimulus-onset. C) The ERP image from subtracting image (B) from (A) is plotted to demonstrate that rERP is effective at removing stimulus and response potentials from trial EEG, as reflected by little consistencies in blue or red hues across trials. D) Grand average residual potentials after “EEG – rERP” (described above) and E) “EEG – ERP” (similar to above description, but substituting standard ERPs for overlap-corrected rERPs) illustrate superiority of rERP approach to standard ERPs in accounting for trial evoked potentials. Note that the overlap-corrected rERP approach better accounts for overlapping stimulus and response potentials because the residual averaged ERP in (D) contains virtually no non-zero potentials while it does in (E).

In **Table 1**, results from logistic regressions reflecting the influence of pMFC-derived Frontocentral P3 and PMP extracted from ERP (top) and rERP (bottom) waveforms on probability of errors are presented. Consistent with the above single-trial analyses, more negative values for Frontocentral P3 and PMP derived from standard ERPs were associated with response errors (*p*_FDR_ < .01, top of **Table 1**). However, when derived from *overlap-corrected rERPs*, only PMP yielded significant associations with errors (*p*_FDR_ < .01) whereas Frontocentral P3 did not (range *p* = .067 to .375), bottom of **Table 1**), indicating that Frontocentral P3’s association with errors may have been confounded by response-locked potentials such as PMP. Here, a one standard deviation larger PMP was associated with approximately 59% *decreased odds* of making an error (i.e., 1 – *OR*, where range *OR* = .41 to .42). Thus, overlap-corrected rERP effects suggest that a smaller PMP (but not Frontocentral P3), is linked to erroneous performance on the trial.

**Table 1.** Prediction of errors using standard ERPs and overlap-corrected rERPs from the pMFC source cluster

| Table 1 |  |  |  |  |  |
| --- | --- | --- | --- | --- | --- |
| Standard trial-averaged ERP |  |  |  |  |  |
| <u>Trial type</u> | <u>Correct</u> | <u>Error</u> | <u>Logistic LMM</u> |  |  |
| ERP measure | *M (SD)* | *M (SD)* | *B<sub>ERROR</sub> (SE)* | *OR* | *p* |
| <u>Congruent</u> |  |  |  |  |  |
| Frontocentral P3 | .33 (.42) | .08 (.64) | -.76 (.21) | .47 | <.001* |
| PMP | .26 (.39) | -.18 (.42) | -1.60 (.30) | .21 | <.001* |
| <u>Incongruent</u> |  |  |  |  |  |
| Frontocentral P3 | .44 (.50) | .05 (.60) | -1.19 (.30) | .30 | <.001* |
| PMP | .41 (.48) | .03 (.51) | -1.10 (.28) | .35 | <.001* |
| Overlap-corrected regression-ERP (rERP) |  |  |  |  |  |
| <u>Trial type</u> | <u>Correct</u> | <u>Error</u> | <u>Logistic LMM</u> |  |  |
| rERP measure | *M (SD)* | *M (SD)* | *B<sub>ERROR</sub> (SE)* | *OR* | *p* |
| <u>Congruent</u> |  |  |  |  |  |
| Frontocentral P3 | .28 (.33) | .18 (1.07) | -.17 (.19) | .84 | .375 |
| PMP | .13 (.31) | -.28 (1.03) | -.90 (.27) | .41 | <.001* |
| <u>Incongruent</u> |  |  |  |  |  |
| Frontocentral P3 | .31 (.43) | .14 (.80) | -.42 (.23) | .66 | .067 |
| PMP | .17 (.36) | -.15 (.81) | -.86 (.31) | .42 | .006* |
| *Note.* Descriptive statistics and results from logistic regressions predicting the log odds probability of an error response per one standard deviation positive change in voltages of standard ERP (top) or overlap-corrected regression ERP measures (rERP, bottom). Positive-valued $B_{ERROR}$ and $OR$s greater than 1 reflect greater odds of an error given greater positivity (larger amplitudes) of Frontocentral P3 (mean value within 340 to 440 milliseconds in stimulus-locked waveforms) or PMP (-160 to -60 milliseconds in response-locked waveforms). We used asterisks to highlight uncorrected $p$-values that survived correction for multiple comparisons at $p_{FDR} < .01$. Other abbreviations: LMM = linear mixed model; SE = standard error; PMP = Pre-Movement Positivity. |  |  |  |  |  |

### Predicting error rates between-subject using factored rERPs

Temporal windows in which pMFC source amplitude in the factored rERP waveforms predicted between-subject differences (*p*_FDR_ < .05) in error rates are depicted by horizontal grey bars in **Figure 11**, as well as waveforms averaged within high (dashed trace) and low (solid trace) error rate tercile groups, reflecting poor and good performers (respectively). Two of these latency regions in the response-locked waveforms temporally corresponded to the PMP window that we found (above) to correlate robustly with errors within the SRI, hereafter referred to as PMP_CORRECT_ (−180 to -78 milliseconds, PMP elicited on congruent correct trials) and PMP_INCONGRUENT_ (−86 to -23 milliseconds, positive deviation in PMP elicited by incongruent stimulus). Mean potentials extracted from these regions were used in remaining mediation analyses.

**Figure 11.**
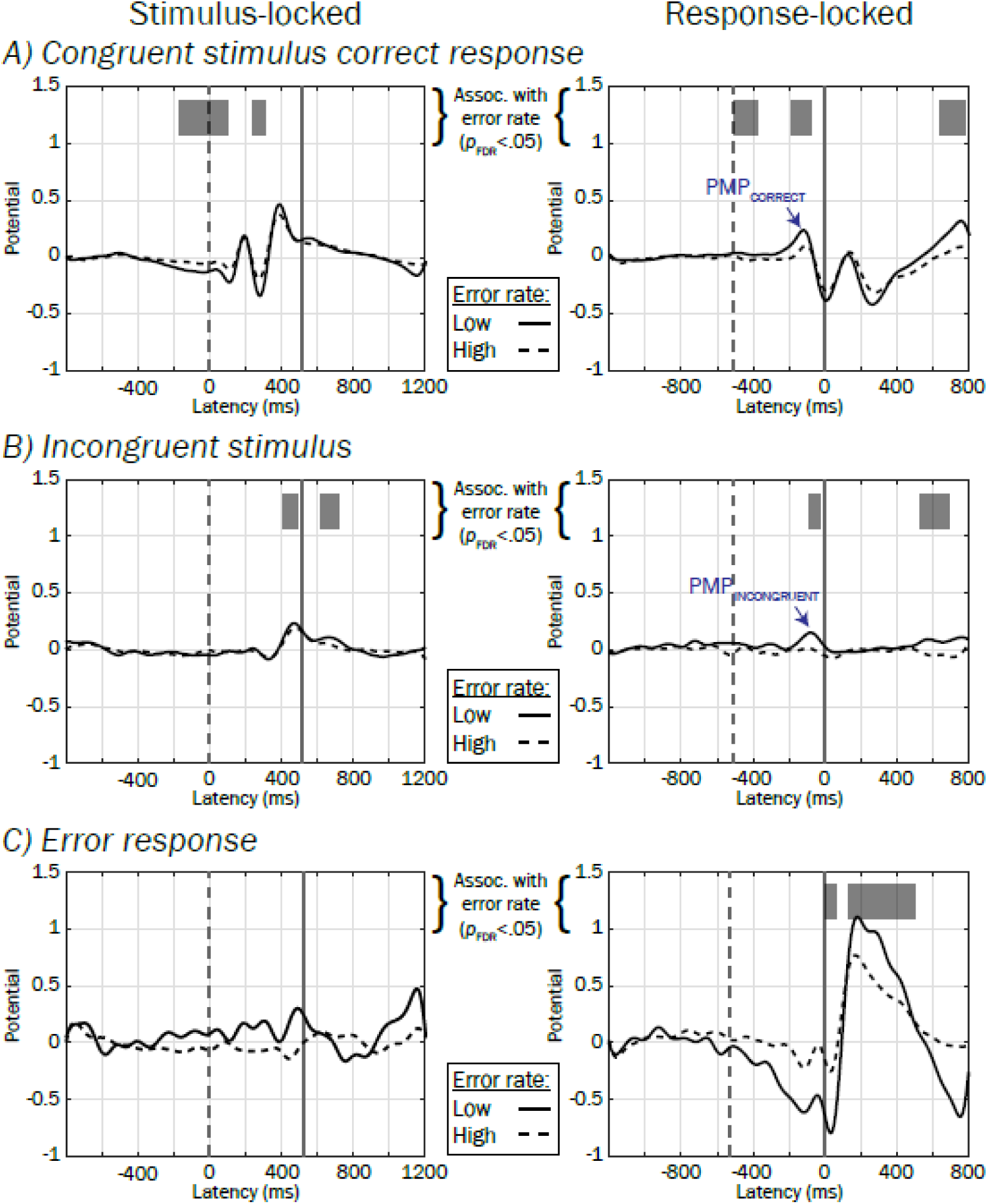
Between-subject prediction of error rates using factored rERPs. Waveforms depicting group averaged factored rERPs within high (dashed traces) and low (solid traces) error rates split by the group median are plotted for illustration. Periods of the factored rERPs are highlighted in greyscale to show regions where there was significant association (*p*_FDR_ < .05 for more than 50 contiguous milliseconds) between rERP and error rate.

### Do amplitude differences in rERPs mediate the association between ADHD symptom counts and heightened task error rates?

ADHD symptoms were associated with elevated error rates on the task (*t*[96] = 3.23, *p* = .002, LMM linear regression), accounting for the main effect of stimulus congruency (incongruent vs. congruent, *t*[137] = 4.50, *p* < .001, LMM linear regression) and the interaction term (incongruent stimulus × ADHD symptoms, *t*[137] = -.29, *p* = .774, LMM linear regression), which was dropped from subsequent analyses. To test whether this main effect of ADHD symptom counts on greater error rates might be mediated by “indirect effects” of the amplitudes of pMFC source potentials PMP_CORRECT_ and PMP_INCONGRUENT_ in the factored rERP (see labeled regions marked with blue arrows in **Figure 11**), we computed paths in **Figure 3B** and presented the results in **Table 2**. Outcomes of mediation tests are shown in the rightmost column and denote whether the confidence intervals for bootstrapped *ab* effects excluded zero, or equivalently, whether the reduction of the “total effect” of ADHD symptoms on error rate (path *c* in **Figure 3A**) to the “direct effect” (path *c’* in **Figure 3B**) was statistically significant.

**Table 2.**
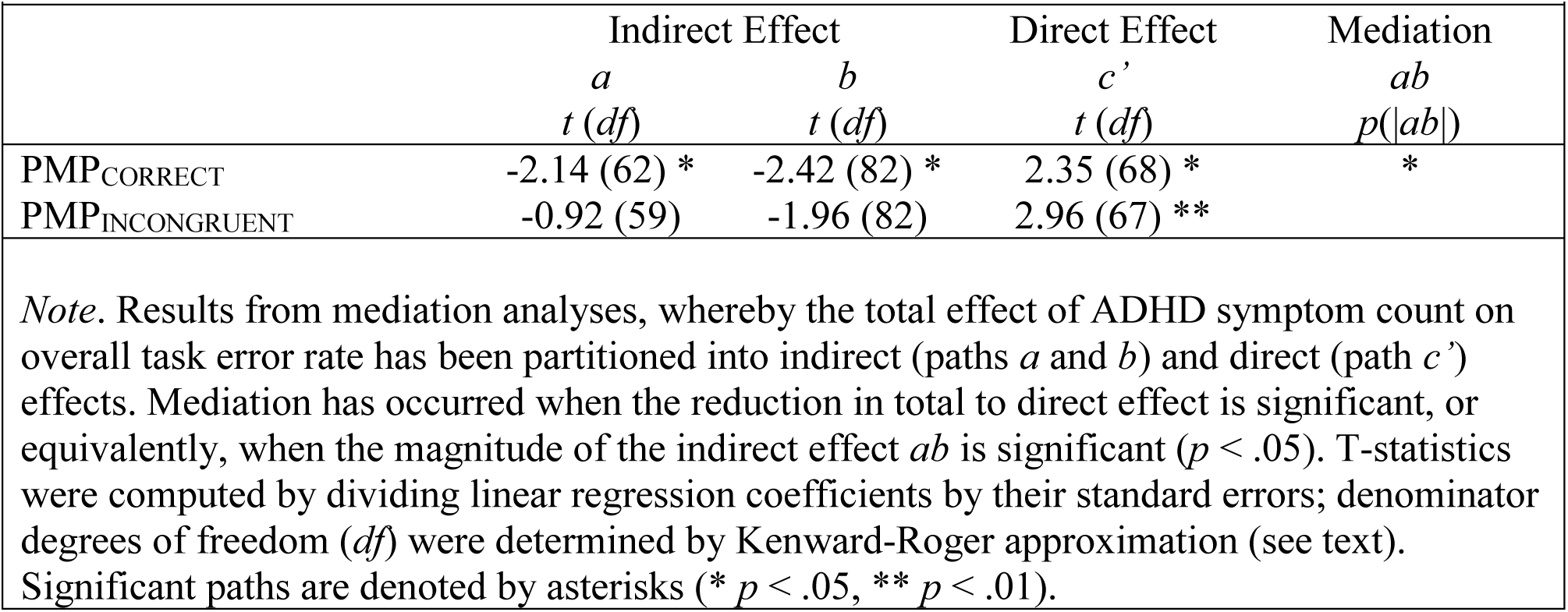
Mediation of ADHD on task error rates using overlap-corrected, factored rERP amplitudes

Only one mediation test was significant, such that including PMPCORRECT amplitude in prediction of error rates alongside ADHD symptoms resulted in a significant reduction of the total effect of ADHD on error rates. Collectively, the negatively-signed *a* path and significant mediation results suggest that reduced pMFC source amplitude in rERPs preceding correct responses during the SRI may partially explain the tendency for adolescents with more ADHD symptoms to make more errors on the task.

## Discussion

We investigated brain potentials in the stimulus-response interval (SRI) during flanker task performance and examined voltage associations with errors, using EEG localized by ICA decomposition and equivalent dipole modeling of brain source processes in a sample of adolescents with varying degrees of ADHD symptoms. We found that smaller Frontocentral P3 and PMP peaks derived from pMFC, which typically occurred within the SRI, predicted errors within subjects. Using a novel regression-ERP (rERP) technique to disambiguate overlapping stimulus- and response-locked brain potentials, the association with errors was attributed primarily to smaller PMP rather than Frontocentral P3, highlighting the importance of response-preceding pMFC processes (PMP) in accurate response selection. We also found a smaller PMP in correct trials to be associated with a larger subject error rate, and that a smaller PMP mediated the strength of the association between ADHD symptom counts and increased error rates.

Our observation of a positive shift in the electrical potential of pMFC sources within 200 milliseconds before the button response in correct performance trials (PMP) is consistent with earlier research on movement-preceding ERPs (Deecke et al., 1969; Makeig et al., 1999; Roman et al., 2005; Delorme et al., 2007). The PMP may reflect a “go signal” (Deecke et al., 1985; Bortoletto et al., 2006) by which supplementary motor areas (pre-SMA and SMA) stimulate a particular motor response to be carried out by circuits involving motor cortex. Indeed, PMP peaked earlier than LRPs projected from putative sensorimotor sources (−110 milliseconds vs. - 75 milliseconds in response-locked ERPs) and we found significant reduction of PMP in pMFC sources on error trials approximately 30 milliseconds before LRPs’ indication of the incorrect motor activation. Thus, our data suggest that *a larger PMP precedes appropriate (vs. inappropriate) motor actions.* A similar effect has been observed for a subset of neurons in monkey pre-SMA which increase firing immediately before correct but not incorrect action selections (Isoda and Hikosaka, 2007). Pre-SMA is thought to facilitate “condition-action associations,” such that conditions (e.g., tasks) require mapping of appropriate actions (here, button responses to target presentations) before action execution (Nachev et al., 2008). Perhaps the PMP generated by pMFC sources before correct responses reflects appropriate condition-action mapping in or near to pre-SMA, whereas on error trials this mapping is incomplete or inappropriate, reflected by a smaller PMP.

We also found that a smaller PMP in correct performance trials was associated with higher subject error rate, and that smaller PMPs mediated the association between subjects’ ADHD symptom counts and higher error rates. To date, PMP has not been investigated with respect to ADHD, although other features of pre-response EEG (e.g., phase variability) have been shown to explain other facets of ADHD performance differences such as RT (e.g., McLoughlin et al., 2013). While it is not known whether the higher frequency of task errors exhibited by ADHD subjects are directly associated with “real-life” accident proneness (e.g., errors in motor vehicle operation; Vaa, 2014), our results suggest that PMP may have clinical implications. For instance, neuromodulation of MFC has improved cognitive performance in healthy individuals (e.g., Spieser et al., 2015); perhaps similar “tuning” of pMFC using PMP as a target for neuromodulation would prove useful in remediating accident proneness in patients with ADHD (e.g., Bloch et al., 2010).

It is important to emphasize that the PMP in our study should not be confused with the “error-preceding positivity” described by other researchers (Ridderinkhof et al., 2003; Allain et al., 2004; Hajcak et al., 2005), which refers to a reduction in the correct-response negativity (Ford, 1999) one *trial* before commencement of the error trial and does not enable insight into action selection brain dynamics within the SRI on the error trial itself (as does PMP). Instead, we believe a related frontocentral scalp-recorded brain potential to the PMP to be the peak preceding the error-related negativity (ERN) noted by others (e.g., Falkenstein et al., 2001; Nieuwenhuis et al., 2001; Debener et al., 2005; Albrecht et al., 2008; Cavanagh et al., 2009; McLoughlin et al., 2009; Albrecht et al., 2010), which has been used (subtracted) for the purpose of quantifying ERN effects but otherwise overlooked. We speculate that PMP might also be conflated with other stimulus-locked ERP peaks that typically precede speeded manual responses such as frontally-focused P2 and P3 waves (e.g., Potts et al., 1996; Makeig et al., 1999; Makeig et al., 2004; Delorme et al., 2007; Perri et al., 2015; Wessel and Aron, 2015). Indeed, in **Table 1** we found the amplitude of Frontocentral P3 to be strongly associated with errors in standard ERPs (*p* < .001), but that these associations were essentially absent after correcting for overlap with response-locked potentials (*p* > .067), suggesting that temporal confounding is perhaps responsible for Frontocentral P3’s seeming association with errors.

## Limitations

It is important to note that because we neither manipulated the size of brain potentials, nor manipulated other factors (e.g., ADHD) influencing task performance, we cannot infer that the size of PMP (or another factor) plays a causal role in determining task accuracy for a given trial or individual. In future research, we speculate that applying neuromodulation (e.g., noninvasive brain stimulation) to pMFC to support PMP amplitude, it may be possible to evaluate such a causal hypothesis.

Consistent with previous ERP studies using flanker tasks (see review by Folstein and Van Petten, 2008), responses were recorded with button presses rather than electromyograms (EMGs), precluding investigation of dynamics accompanying central motor conductance timing (CMCTs, delays between motor brain potentials and EMG responses) and EMG onset-to-button press timing (e.g., “partial errors"; Roger et al., 2014). While we cannot know how CMCT- and EMG-induced dynamics might influence the latencies of brain potentials in our study, we conjecture that such effects are small (e.g., CMCTs typically range 3 to 5 milliseconds in humans aged > 4 years; Udupa and Chen, 2013) or relatively constant (i.e., shifting waveforms’ latencies by some fixed delay) given the tight succession of EMG and button presses. Crucially, we observed PMP concurrently with well-known ERP features, such as LRPs, that typically precede EMG responses. To the extent that PMP effects accompany or precede LRPs, it may be assumed that they also tend to precede EMG onsets.

The spread of brain locations of the equivalent dipoles in the pMFC source cluster appears larger than expected with a single functional brain region (see Tsai et al., 2014). Dipole localization errors may have added to this spread; these could arise from insufficiencies in subject head models (e.g., inaccurate skull conductivity; Akalin Acar and Makeig, 2013) or in co-registration of subject dipoles to the MNI template head. Skull conductivity varies across individuals due to several factors (e.g., development) but was held constant here across subjects, as a noninvasive method for its estimation within-subject (Akalin Acar et al., 2016) has not yet been made readily available. Lastly, pMFC source clusters originally obtained by the k-means algorithm did not include all subjects, necessitating post-hoc addition of outlier sources from k-means unincluded subjects for the sake of preserving sample size and statistical power. Although this step may introduce possible inhomogeneities in the clusters themselves, added sources appeared comparable to original sources (e.g., **Figure 5**).

## Conclusions

Using source-localized EEGs resolved by ICA decomposition and subject-specific head models, we found that smaller Frontocentral P3 and PMP amplitudes projected from pMFC, typically occurring within the SRI, were associated with error commission in a flanker task. After regressing out stimulus-from response-locked contributions using a novel regression-ERP technique, errors appeared largely dependent upon response-preceding (“action selection,” PMP) processes, instead of stimulus-evoked (“perceptual,” Frontocentral P3) processes. Finally, having more ADHD symptoms was associated with producing higher error rates and smaller PMP partially mediated this association.

We propose that reduced PMP could be a valuable target for intervention as well as being possibly useful as a developmental endophenotypic biomarker reflecting genetic risk for error proneness in psychopathology including ADHD (cf. Iacono and Malone, 2011; Burwell et al., 2016; Iacono et al., 2016). Indeed, PMP amplitudes in response-locked rERPs from correct trials were similar among identical twin pairs (intra-class correlation = .29, *p* = .023), reflecting familial influences. The putative genetic basis of PMP remains to be investigated, but it is possible that the source decomposition and/or rERP methodologies used in our study could aide in clarifying the degree to which brain potentials reliably tap into genetic mechanisms underlying psychopathology, which may have been otherwise confounded in scalp EEG recordings and standard ERPs (McLoughlin et al., 2014; Loo et al., 2016). Regardless, these results bolster the importance of pMFC during the SRI and suggest a role in correct and erroneous action selections. We suggest that future research should both further our understanding of PMP and determine the degree to which reduced PMP may be involved in the etiology of error proneness in psychopathology such as ADHD.

## Acknowledgements

This research was made possible by funding from grant number AA017314 from the National Institute of Alcohol Abuse and Alcoholism (NIAAA) as well as grant numbers DA036216 and DA05147 from the National Institute on Drug Abuse (NIDA). In addition, SJB is funded by a NIDA T32 postdoctoral fellowship (DA037183) and received funding from the Society for Psychophysiological Research that enabled mentorship from SM at the Swartz Center for Computational Neuroscience (SCCN) at UCSD. SJB would also like to thank Dr. Zeynep Akalin Acar and Matthew Burns at the SCCN for providing feedback in computational head modeling and regression ERP procedures (respectively), as well as Dr. Philip Burton at UMN’s Center for Magnetic Resonance Research for his expert guidance in the cross-subject dipole co-registration method. SM’s participation was funded by a gift from The Swartz Foundation (Old Field, NY).

